# Revisiting reptile home ranges: moving beyond traditional estimators with dynamic Brownian Bridge Movement Models

**DOI:** 10.1101/2020.02.10.941278

**Authors:** Inês Silva, Matt Crane, Benjamin Michael Marshall, Colin Thomas Strine

## Abstract

Animal movement, expressed through home ranges, can offer insights into spatial and habitat requirements. However, home range estimation methods vary, directly impacting conclusions. Recent technological advances in animal tracking (GPS and satellite tags), have enabled new methods for home range estimation, but so far have primarily targeted mammal and avian movement patterns. Most reptile home range studies only make use of two older estimation methods: Minimum Convex Polygons (MCP) and Kernel Density Estimators (KDE), particularly with the Least Squares Cross Validation (LSCV) and reference (*h*_*ref*_) bandwidth selection algorithms. The unique characteristics of reptile movement patterns (*e.g.* low movement frequency, long stop-over periods), prompt an investigation into whether newer movement-based methods –such as dynamic Brownian Bridge Movement Models (dBBMMs)– are applicable to Very High Frequency (VHF) radio-telemetry tracking data. To assess home range estimation methods for reptile telemetry data, we simulated animal movement data for three archetypical reptile species: a highly mobile active hunter, an ambush predator with long-distance moves and long-term sheltering periods, and an ambush predator with short-distance moves and short-term sheltering periods. We compared traditionally used home range estimators, MCP and KDE, with dBBMMs, across eight feasible VHF field sampling regimes for reptiles, varying from one data point every four daylight hours, to once per month. Although originally designed for GPS tracking studies, we found that dBBMMs outperformed MCPs and KDE *h*_ref_ across all tracking regimes, with only KDE LSCV performing comparably at some higher-frequency sampling regimes. The performance of the LSCV algorithm significantly declined with lower-tracking-frequency regimes, whereas dBBMMs error rates remained more stable. We recommend dBBMMs as a viable alternative to MCP and KDE methods for reptile VHF telemetry data: it works under contemporary tracking protocols and provides more stable estimates, improving comparisons across regimes, individuals and species.

## 1. Introduction

Animal movement is an underlying process in many ecological systems, and there is a growing understanding of how individuals behave through space and time (Nathan *et al.*, 2008; Gurarie *et al.*, 2016). Movement is often conceptualized then presented as a home range, defined as the area animals move through during “normal” activities, including resource acquisition and reproduction (Burt, 1943; Powell 2012). While the utility of the home range concept has been questioned in recent years (Kie *et al.*, 2010; Powell & Mitchell, 2012), home range estimates continue to have a range of applications: identifying behavioural adaptations to predictable environmental features (Riotte-Lambert & Matthiopoulos, 2019) or inferring habitat use (Fisher, 2000; Dickson & Beier, 2002; Tikkanen *et al.*, 2018; Marshall *et al.*, 2019). Applying a home range approach to ecological research questions requires careful consideration (Péron, 2019), as any conclusions drawn can be profoundly impacted by the natural history of the target species.

Terrestrial reptiles —broadly lizards, snakes, and tortoises— have *distinct* natural histories from mammals (*e.g.* as ectotherms), resulting in *distinct* movement patterns. Many reptiles move less frequently than comparatively sized mammals (Hailey, 1989), but more importantly, many terrestrial reptiles spend prolonged periods stationary under shelter (one day to several weeks; Guarino, 2002; Bruton, McAlpine, Smith, & Franklin, 2014; Mata-Silva, DeSantis, Wagler, & Johnson, 2018). These inconsistent movement patterns severely impact inferences drawn from home range analyses.

To properly inform desperately needed conservation actions (Gibbons *et al.*, 2000; Roll *et al.*, 2017), we must tailor our methodologies to the peculiarities of reptile movement (Péron, 2019) –otherwise we risk designing suboptimal solutions. We must assess the utility of newer methods designed for mammals, before applying them to reptiles (Silva, Crane, Suwanwaree, Strine, Goode, 2018).

With the rise of Global Positioning System (GPS) animal tracking, researchers have developed new statistical approaches for calculating home ranges that take advantage of the high number of location fixes. However, GPS tracking currently has limited application in reptiles (see Schofield *et al.*, 2007; Campbell *et al.*, 2013; Rosenblatt *et al.*, 2013; Smith, Hart, Mazzotti, Basille, & Romagosa, 2018) as their natural history poses several problems (Hebblewhite & Haydon, 2010; Wolfe, Fleming, & Bateman, 2018); *e.g.* weakened signal due to the surgical implantation or attachment of the tag, limited number of species which can be ethically attached due to body size (Smith *et al.*, 2018), reduced fix rate and precision due to sheltering underground (Bruton *et al.*, 2014, Wolfe *et al.*, 2018).

Given that traditional home range estimators –Minimum Convex Polygons (MCP) and Kernel Density Estimators (KDE)– present major limitations for telemetry-based reptile studies (see Row & Blouin-Demers, 2006), it is important to investigate whether newer methods developed for GPS tracking data can be applied to reptile-targeted Very High Frequency (VHF) radio-telemetry studies. Dynamic Brownian Bridge Movement Models (dBBMMs) are a technique intended for GPS telemetry, allowing for efficient and repeatable analysis of high-resolution data –particularly useful for animals with behaviourally distinct movement patterns. The method creates a one-dimensional fix-frequency independent behavioural measure (Brownian motion variance; Kranstauber, Kays, LaPoint, Wikelski, & Safi, 2012) that have been employed to elucidate avian and mammal home range and movement patterns (*e.g*. Palm *et al*., 2015; Byrne, McCoy, Hinton, Chamberlain, & Collier, 2014; Lai, Bêty, & Berteaux, 2015; Buechley, McGrady, Çoban, & Sekercioglu, 2018).

Leveraging dBBMMs may benefit VHF studies (Silva *et al.*, 2018; Walter, Onorato, & Fischer, 2015); and while multiple simulations studies have investigated how different methods interact with animal movement and home range delineation (*e.g*. Katajisto & Moilanen, 2006; Row & Blouin-Demers, 2006; Knight *et al*., 2009; Cohen, Prebyl, Collier, & Chamberlain, 2018), none have targeted reptile-specific movement patterns.

We assess home range estimates resulting from variable study designs common in the reptile spatial ecology literature: namely temporally low-resolution tracking regimes. We simulate movement data of three archetypal reptile species, thoroughly examining the most common home range estimators — Minimum Convex Polygons (MCPs) and Kernel Density Estimators (KDEs). We next compare traditional estimators to a newer method: dynamic Brownian Bridge Movement Models (dBBMMs). Finally, we discuss the implications of home range estimator choice, and present guiding principles for reptile spatial ecology sampling designs.

## 2. Materials and Methods

### 2.1. Simulated animal movement and tracking data

We used the *SimData* function in the *momentuHMM* package (McClintock & Michelot, 2018) to simulate movement data from a Hidden Markov Model (HMM). HMMs are time-series models where the movement pattern of an animal is assumed to depend on the underlying behavioural state of the animal (Langrock *et al.*, 2014). We simulated data for 32 individuals from three archetype reptile species, to represent three main groups within reptile movement ecology: **Species 1** corresponds to highly mobile (active hunters) with long-term shelter sites (*e.g.* monitor lizards, some skinks, and elapids like mambas and king cobras); **Species 2** represents less mobile reptiles, capable of moving long distances but are ambush foragers, and will still shelter for long periods (*e.g.* pythons); finally, **Species 3** represents smaller ambush predators, infrequently moving and sheltering for shorter periods (*e.g.* viperid snakes, some smaller lizard species).

Each archetype had a unique set of state-dependent parameters and transition probabilities with the same three behaviour states: “sheltering” (state 1), “moving” (state 2), “resting” (state 3). The state-dependent data streams included step length (l_*t*_) and turning angle (θ_*t*_), which we generated from Gamma and von Mises distributions, respectively. The simulations included a spatially correlated covariate for state 2, to reflect habitat preferences, while states 1 and 3 followed a *cosinor* function, to reflect cyclical patterns of long-term sheltering (state 1) and circadian rhythms (state 3). To simulate individual variation and movement in a heterogeneous landscape we generated a random neutral landscape with fractal Brownian movement, using the *NLMR* package (Sciaini, Fritsch, Scherer, & Simpkins, 2018). For further details on these simulated species, as well as their specific step lengths, turning angles and transitional probabilities, see Appendix S1, Supporting Information.

After creating the full simulated data set (regime 1), we generated six subsets of the data to represent various field sampling regimes (regime 2-7): four locations per day, two locations per day, one location per day, two locations per week, one location per week, and one location per month (Figure 1). For each subset, we assumed a consistent regularly scheduled sampling protocol limited to the species’ activity periods.

**Figure 1.**
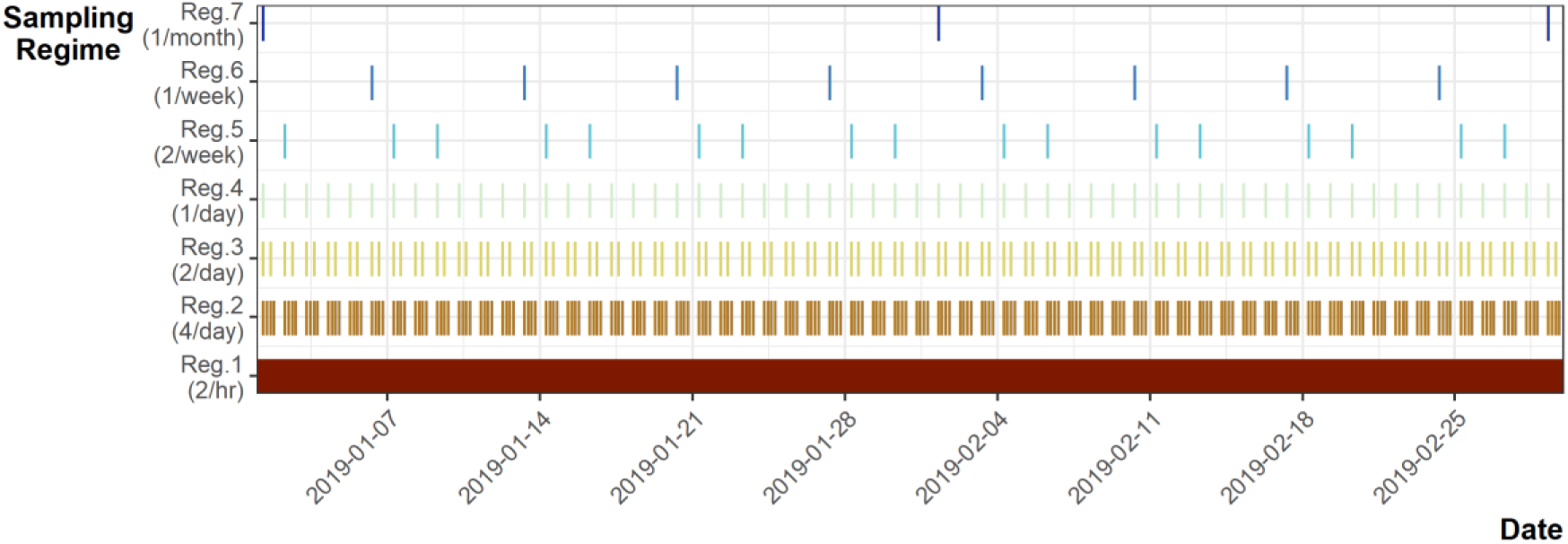
Example two-month period showing how data is thinned to represent different tracking regimes.

The autocorrelated nature of tracking data poses difficulties for home range estimators that assume independence between points, namely KDEs. Attempting to remove autocorrelation to fit these assumptions can reduce the biological relevance of the home range (De Solla *et al*., 1999), but advocated in reptile home range studies (Swihart & Slade, 1985; Worton, 1987).

We investigated the temporal autocorrelation present in our simulated dataset to determine whether our coarser sampling regimes compiled with KDE independence assumptions. Other than less frequent tracking, autocorrelation may be reduced by removing repeated locations. This method is particularly relevant for reptiles that exhibit long term sheltering. We considered this special case –sampling regime 8– where only animal relocations are included in the home range estimation. For regime 8 we used the four location per day sampling regime, and then removed data points where the animal was stationary.

We described the autocorrelation in the simulated data using the *ctmm* package’s variogram functionality (Calabrese, Fleming, & Gurarie, 2016; Fleming *et al.*, 2017), and plotted the minimum number of days until the autocorrelation became insignificant with raincloud plot code from Allen, Poggiali, Whitaker, Marshall, & Kievit (2019).

### 2.2. Home range estimators

#### 2.2.1. Minimum convex polygon

We calculated the Minimum Convex Polygon (MCP) for each simulated individual that created the smallest area convex polygon containing all animal locations. We used the 95% MCP, which removes outlying points on the assumption that these represent exploratory movements and thus not part of the home range, as originally defined by Burt (1943). The MCP method has long been lauded as a way of maintaining comparability and historical consistency with previous studies (Jennrich & Turner, 1969), yet has well documented issues: extreme sensitivity to sampling size and tracking duration (Anderson, 1982), and overestimated boundary delineation (Robertson, Aebischer, Kenwards, Hanski, & Williams, 1998), with the inclusion of areas that the animals never use (Börger *et al*., 2006; Laver & Kelly, 2008). However, Row & Blouin-Demers (2006) argued that MCPs are preferable to kernel density estimators specifically for herpetofauna, and MCPs’ use persists for “comparisons” in reptile telemetry studies (Petersen, Goetz, Dreslik, Kleopfer, & Savitzky, 2019). An additional and considerable limitation of MCPs is that they do not create a probabilistic utilization distribution.

#### 2.2.2. Fixed kernel home range

Fixed KDE home ranges rely on a smoothing parameter (bandwidth) to generate a utilization distribution. Bandwidth selection for KDE can dramatically influence home range estimation (Seaman *et al.*, 1999), and thus we included two bandwidth selection algorithms, reference bandwidth (*h*_*ref*_) and Least-Squares Cross-Validation (LSCV), for our comparisons. Both bandwidth selection methods are frequently used in reptile VHF studies, but potentially flawed for herpetofauna (Row & Blouin-Demer, 2006). The *h*_*ref*_ method tends to overestimate home ranges while LSCV tends to underestimate (Hemson *et al.*, 2005). In general, fixed KDE home ranges are not accurate when using autocorrelated data regardless of bandwidth selection function (Noonan *et al*., 2018).

#### 2.2.3. Dynamic Brownian Bridge Movement Model

Dynamic Brownian Bridge Movement Models (dBBMMs) provide utilization distributions based on animal movement paths. The method accounts for temporal autocorrelation, so it requires all locations to be time stamped. In addition, dBBMMs incorporate error associated with each triangulated location, which we kept consistent across species and regimes (at 5 metres) for the following reasons: (1) neither MPCs nor KDEs account for location error, so the evaluation of the impact of this metric would be solely on one method and not effective for comparison purposes; (2) location error associated with VHF telemetry is extremely variable, dependent on macro and micro-habitat characteristics as well as tracking protocols (which we are not assessing); and (3) we wanted to account for cases where GPS error can be greater than step length (e.g. viperids, small lizards). The dBBMM method also allows calculation of Brownian motion variance (*σ*^*2*^*m*), which can help researchers determine how movement trajectories can occur due to a species’ behaviour and activity (Kranstauber *et al.*, 2012). Motion variance can help detect breeding and foraging behaviour in reptiles, even with VHF telemetry data (Silva *et al.*, 2018).

### 2.3 Method comparison

To compare the error generated from each home range estimator, we calculated the overlap with the theoretical “true home range” for each individual. We generated an individual’s “true home range” by creating a buffer around all the simulated movement points with a width of two-times the step length intersect from each simulated species’ movement state (40-m for Species 1, 20-m for Species 2, 10-m for Species 3). This provided a conservative home range estimate (excluding the impact of habitat), but more generous and biologically sensible than only using simulated movement pathways. For each home range, we calculated the omission (Type I, false positive) and commission (Type II error, false negative), using the 95% contours for MCP, KDE and dBBMMs. We used the 95% contours, as this is the standard level used in most home range estimates. We then calculated the F-measure [2/(recall^-1^+precision^-1^)], which provides a balanced metric of Type I and Type II errors and is insensitive to true negative rates (Sofaer, Hoeting, & Jarnevich, 2019).

We explored the relationship between methods, regimes, and F-measures using a Bayesian generalized linear mixed model with the *brms* package (Bürkner, 2017). We specified a model set for each species, with F-measure as our response variable following a beta distribution (as it is bound between 0 and 1), with individual as a random effect to account for individual variation and a varying slope for the effect of method. We excluded regime 8 (four locations a day, relocations only) as this sampling regime was not systematic. We ran models with six Markov Chain Monte Carlo (MCMC) chains, each with 6,000 iterations (1,000 burn-in iterations, thin = 1), and we set Δ to 0.99. We fitted each model with half-Cauchy weakly informative priors (Lemoine, 2019). We checked model convergence by inspecting trace plots and 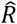 values (Bürkner, 2017), assessed model fit visually via posterior predictive diagnostic plots, and evaluated model performance using leave-one-out cross-validation (Vehtari *et al*., 2017) and Bayesian *R*^*2*^. For further details on model selection and validation, see Appendix S2, Supporting Information.

We compared the special case of regime 8 (similar to regime 2 but only relocation points) to the original regime 2 in its own Bayesian model set; this allowed us to evaluate the impact of removing stationary locations as a method of reducing data autocorrelation. Additionally, for this special case we only compared the best performing KDE bandwidth (LSCV) and dBBMMs.

All datasets and R code to reproduce analyses is available at Zenodo repository platform (DOI:10.5281/zenodo.3660796). We wrote code for R (v.3.5.2, R Core Team), using R studio (v.1.2.1335, R Studio Team).

## 3. Results

### 3.1. Simulated animal movement and tracking data

The complete dataset for each simulated individual consisted of *n* = 17,521 data points for a full year, with 30-minute time steps (regime 1). Each regime progressively lowered the available data (*n*^reg 2^ = 1,460 data points, *n*^eg 3^ = 730, *n*^reg 4^ = 365, *n*^reg 5^ = 104, *n*^reg 6^ = 52, *n*^reg 7^ = 12), while regime 8 varied for each species and individual due to the variability in sheltering and resting behaviour (*n*^species 1^ *=* 5,189 ± 204 data points (mean ± SD); *n*^species 2^ *=* 3,501 ± 1,099; *n*^species 3^ *=* 3,873 ± 573). Visual validation of movement patterns matched with reported patterns in the literature (*e.g.* Parent & Weatherhead, 2000; Reed & Douglas, 2002; Wasko & Sasa, 2009; Hart *et al*., 2015; Smith *et al.*, 2018; Silva *et al*., 2018; Marshall *et al.*, 2019), and the predicted patterns of the three archetypes (Figure 2).

**Figure 2.**
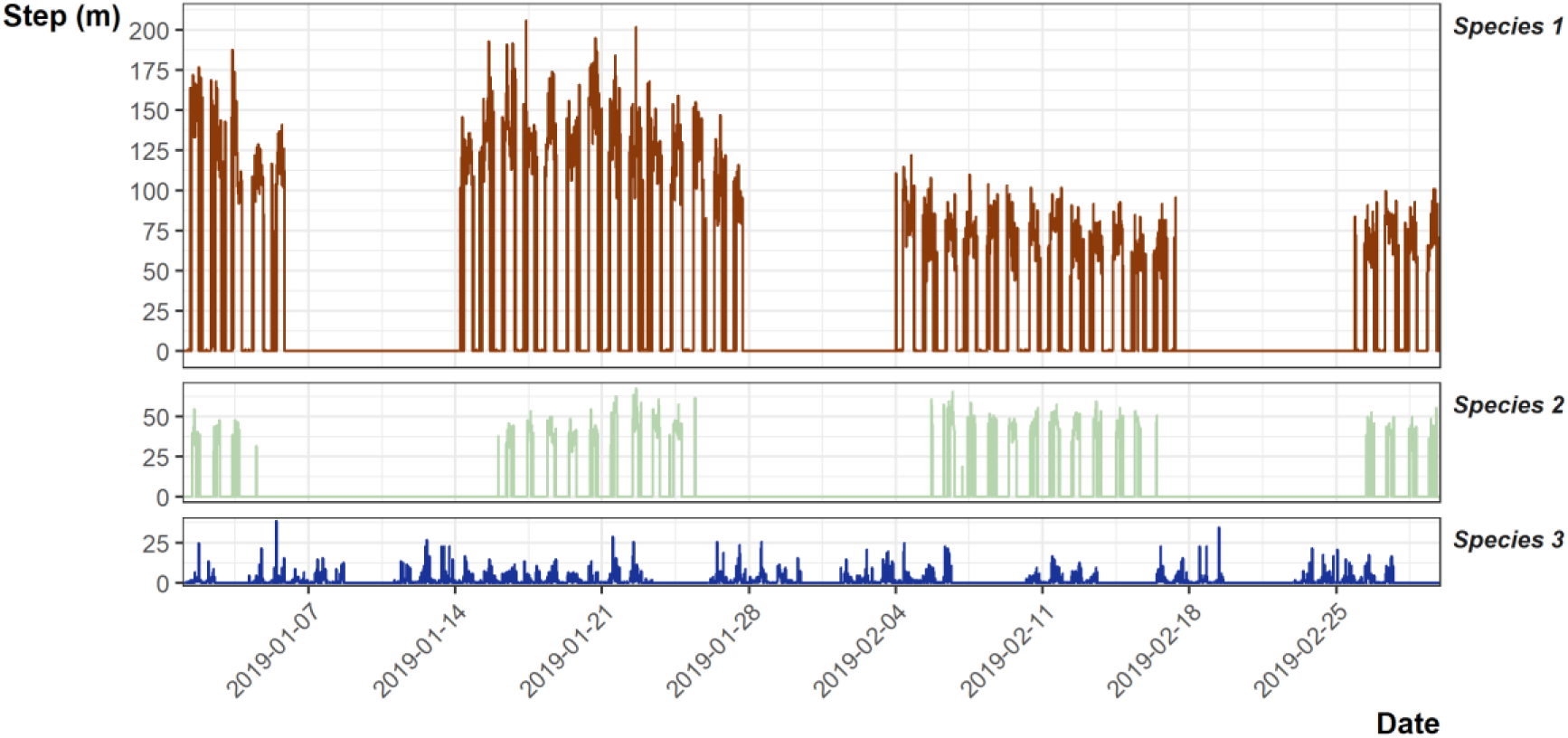
Example two-month period illustrating how step distance (m) and its frequency differs between our three species archetypes.

As expected, all simulated species and individual datasets showed strong autocorrelated structure. Time until insignificant autocorrelation far exceeded even the coarsest tracking regime tested (regime 7, *i.e.* 1/month), indicating that all tracking regimes breach the assumption of independence required for KDE methods (Figure 3).

**Figure 3.**
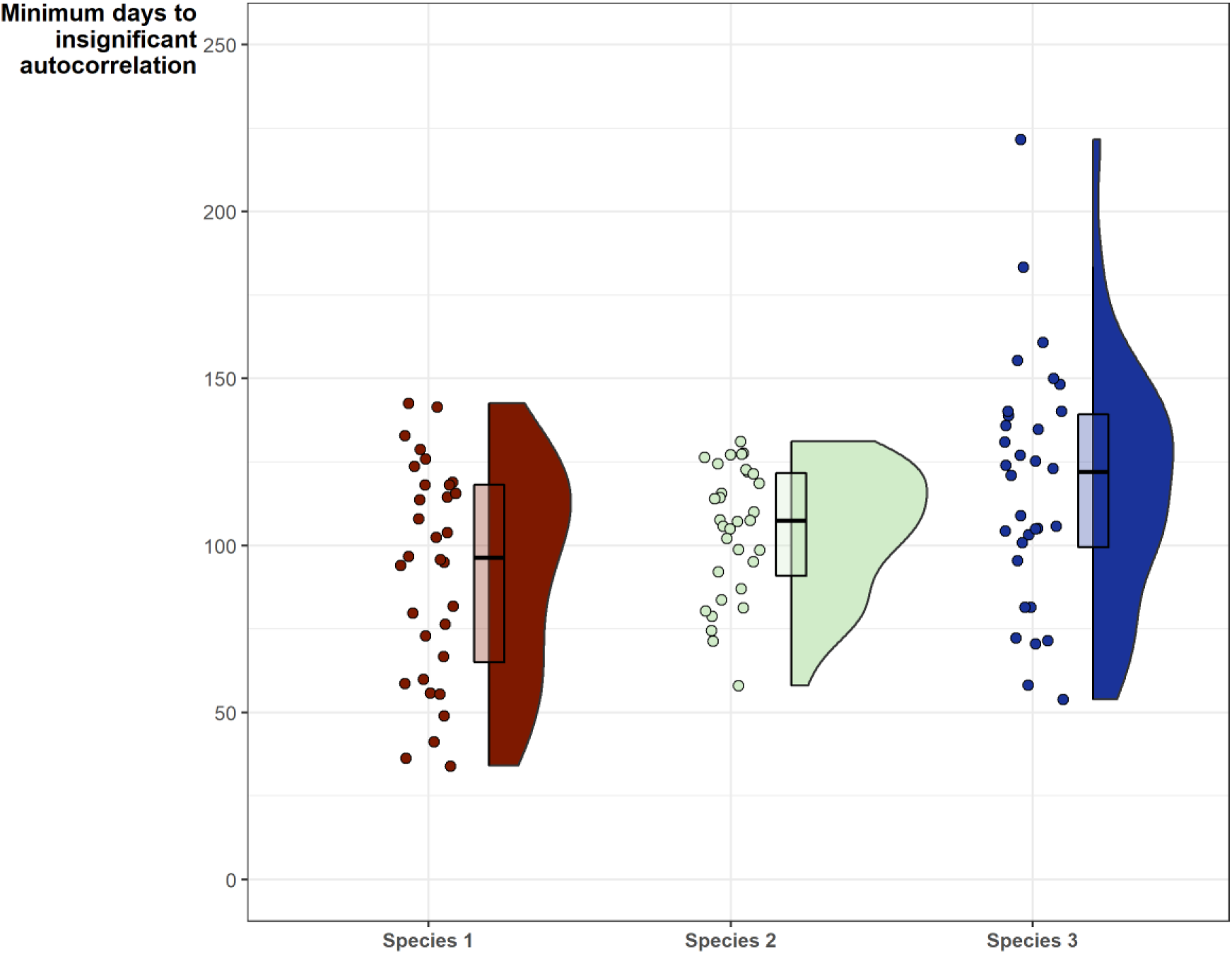
Minimum number of sampling days until the autocorrelation becomes insignificant and data points can be considered independent.

### 3.2. Method comparison: Omission vs. commission

Overall, coarser tracking regimes lead to greater % error when compared to true home ranges. However, the balance between omission and commission is inconsistent and varies between home range estimation methods (Figure 4). There is also a general trend towards commission error when estimating home ranges because omission error is bounded between 0 and 100%.

**Figure 4.**
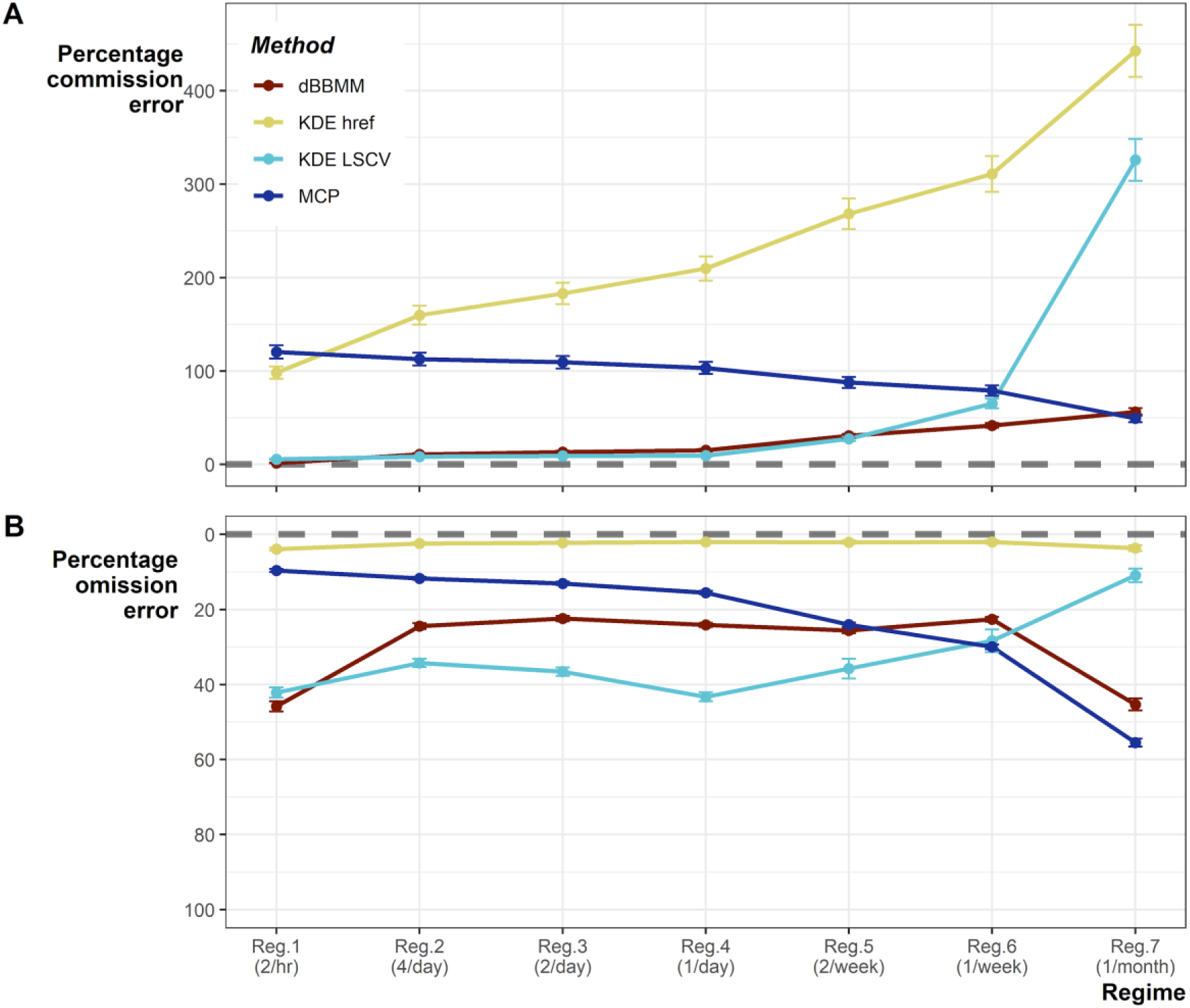
Percentage error from the true home range using 95% contours. **A)** Commission error represented by positive values **B)** omission error represented by negative values. Error bars represent standard error of means across species (3) and individuals (96). Note the different scales for error, as omission error cannot exceed 100% of the true home range area.

#### 3.2.1. Minimum convex polygon

Minimum convex polygons were the only method that showed a constant offset between omission and commission, as one increases the other decreases nearly 1:1. In addition, MCPs were the only method that decreased their commission error as tracking regime became temporally coarser. At frequent tracking regimes, MCPs only introduced minimal omission error, but their starkest failure is in their simple shape leading to the greatest commission error at highest resolution tracking regime (Figure 4, 5).

**Figure 5.**
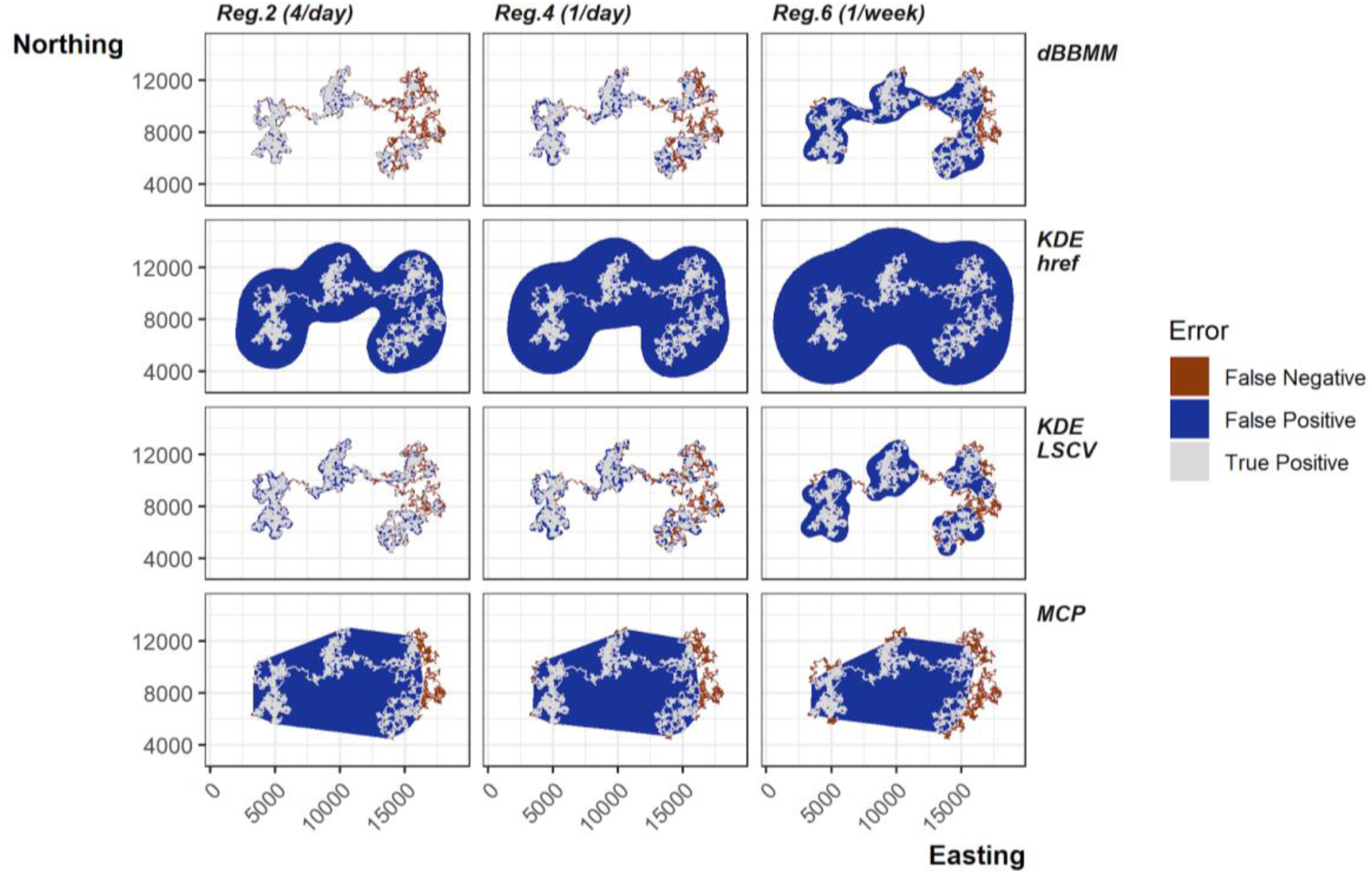
An example of how method and regime can interact to produce different levels of false negative, false positive against the true home range. All contours shown are produced from the 95% utilization distribution.

#### 3.2.2. Fixed kernel home range

The fixed kernel home range using *h*_*ref*_ smoothing factor was by far the worst estimator for commission error. At low resolution tracking regimes, the >400% overestimation leads to near complete loss of home range edge fidelity (Figure 4). Due to this heavy emphasis on generous home range estimation KDE *h*_*ref*_ produced negligible omission error (Figure 5).

By comparison KDE LSCV produced consistently lower commission error at higher resolution tracking regimes, but once the regime was once a week or coarser LSCV commission error spikes (up to 300% overestimation). LSCV consistently performed worse in terms of omission error when applied to tracking regimes with multiple tracks per day. Additionally, the LSCV algorithm frequently failed to converge (68.5% of all LSCV home ranges failed). Only regime 7 converged consistently; the inclusion of more data exacerbated convergence failure (regime 1-4, 100%; regime 5, 43.8%; regime 6, 33.3%). Using only relocations reduced convergence failures (regime 8, 2.08%) compared to its closest parallel regime (regime 2,100%).

For both KDE methods, omission and commission error variability (displayed as SE on Figure 4) increased as tracking regime became coarser.

#### 3.2.3. Dynamic Brownian bridge movement model

Overall dBBMMs performed best. The method produced low commission error levels, matching KDE LSCV performance (Figure 4). Unlike LSCV, dBBMMs commission error remained more stable and lower when applied to coarser tracking regimes. Only MCPs produced a comparative level of commission error at the coarsest tracking regimes, but dBBMMs kept some semblance of shape fidelity and connectivity (Figure 5). Unlike other methods, dBBMM error remained low and balanced between omission and commission, never exceeding 75%.

#### 3.2.4. Special case of regime 8

Tracking regime 8 (four locations per day, relocations only) cannot be directly compared to the other regimes as the structure of the tracking is different. A fairer comparison is between regime 8 and 2 (four locations per day). Similar to all other regimes, regime 8 fails to remove autocorrelation to insignificance (Figure 3); however, it did improve the performance of KDE LSCV estimation despite still breaching the fundamental independence assumption (Figure 5, 6). The removal of repeated stationary points prevented the LSCV smoothing from grouping too tightly to point concentrations (i.e. long-term shelter sites), ultimately countering the tendency towards omission error for LSCV. However, on average, dBBMMs performed very similarly and balanced the omission and commission well (Figure 4). The dBBMMs had the added advantage of assuming serial dependence of points and, unlike LSCV, perform well when provided high quantities of data.

**Figure 6.**
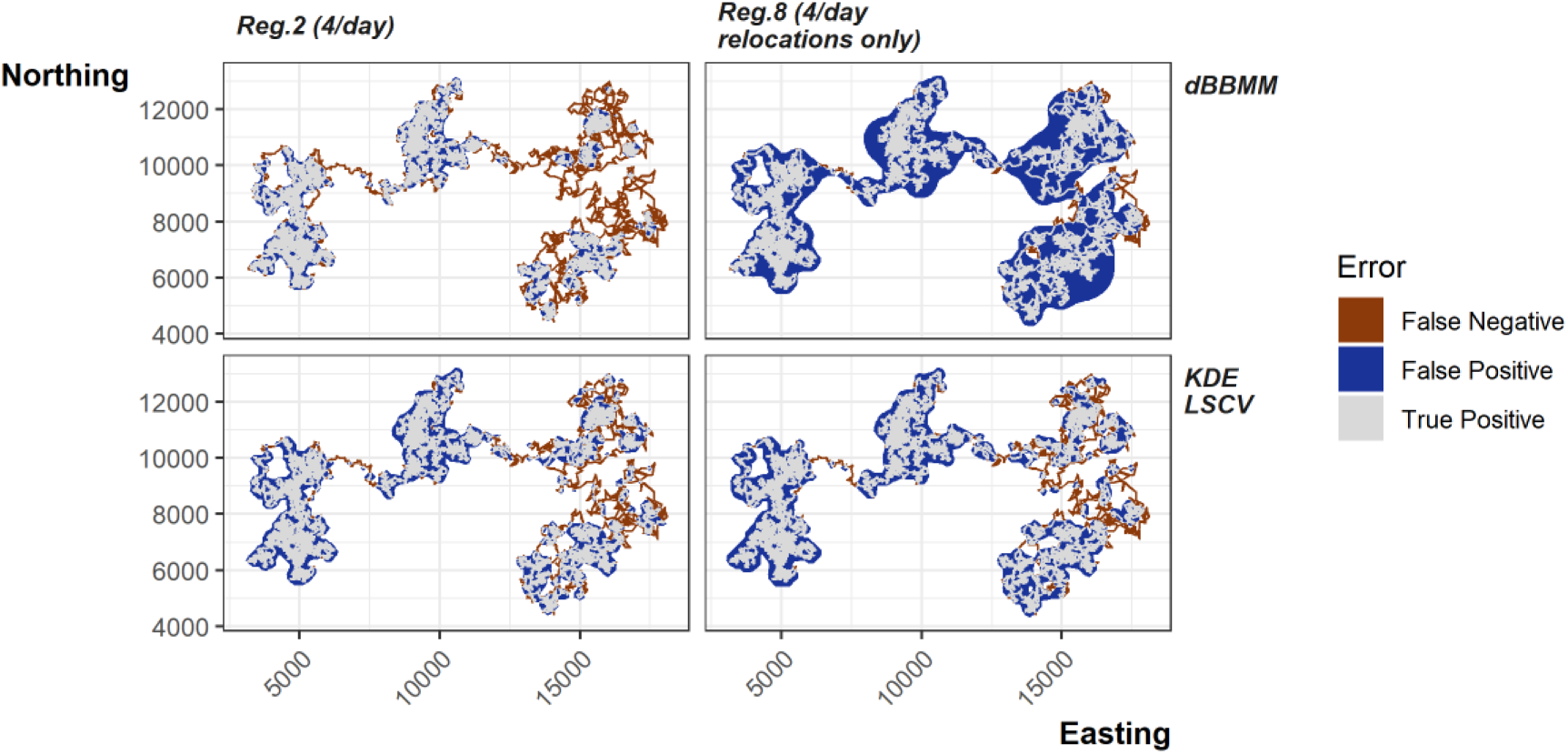
Comparison between the error rates produced by the KDE LSCV and dBBMM 95% contour ranges when using data from sampling regime 2 (four locations per day) and regime 8 (four locations per day, relocations only).

**Figure 7.**
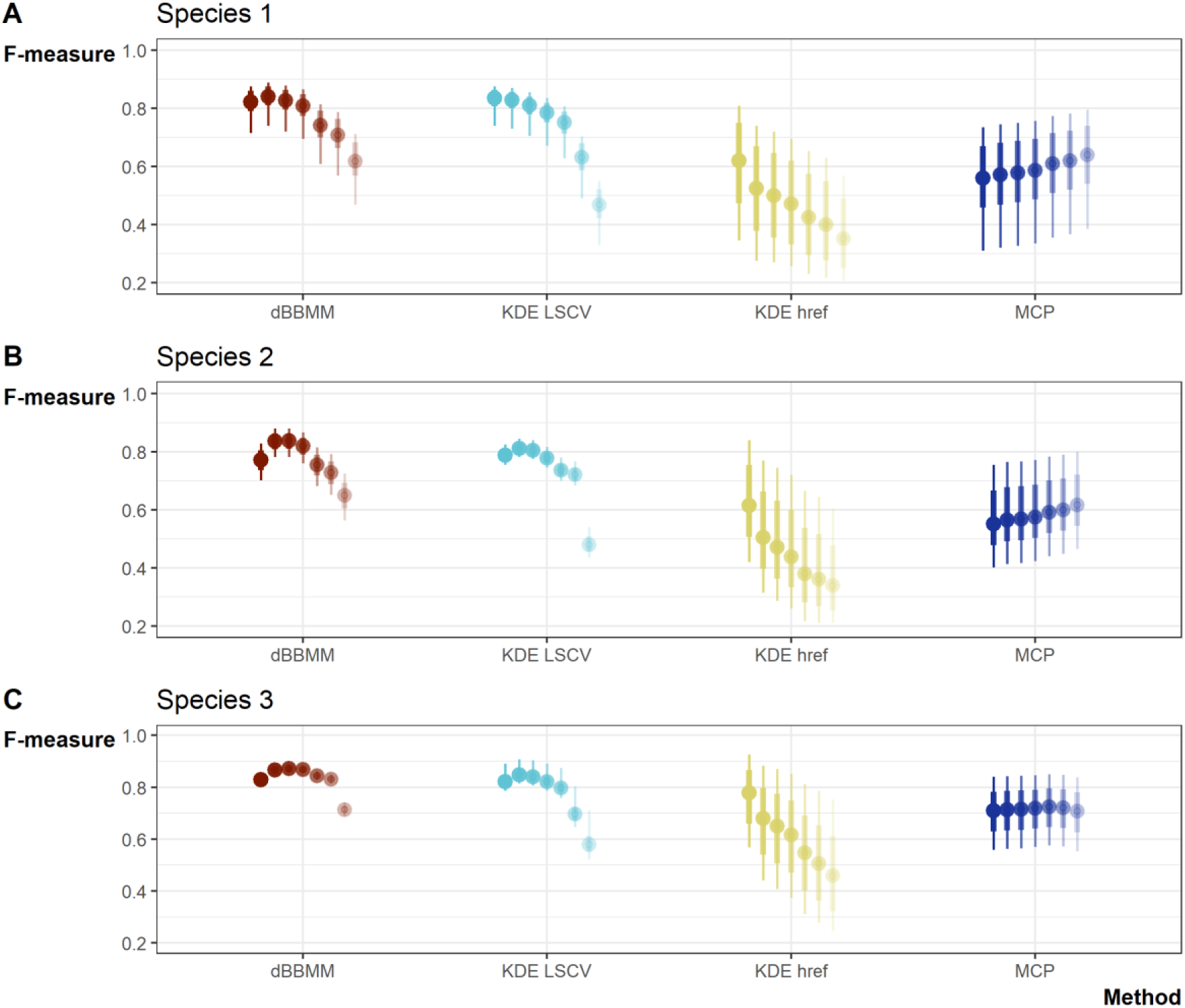
Model results that aimed to predict F-measures using method, regime, and individual ID by species. Tracking regime 1-7 are shown left to right with lowering levels of opacity. Fitted draws were taken only from the first 5000 samples.

### 3.3. Method comparison: F-measures

The Bayesian models converged and performed well for all three species, with 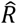 values ≃ 1.00 (Appendix S2, Supporting Information), and *R*^*2*^ values indicating considerable predictive power (Species 1: Bayesian *R*^*2*^ = 0.960, 95% CrI: 0.958–0.962; Species 2: Bayesian *R*^*2*^ = 0.946, CrI: 0.755–0.786; Species 3: Bayesian *R*^*2*^ = 0.905, CrI: 0.897–0.911). Overall, our best models showed an interaction effect of methods and regimes on F-measures; all species had a non-zero positive relationship between F-measures and regimes, with higher estimates for dBBMM and KDE LSCV, while both MCP and KDE *h*_*ref*_ showed considerably worse F-measures. However, Species 1 home range estimations were associated with lower F-measures, suggesting that the home ranges of species with high movement and long periods of sheltering are harder to model than those with more stable movement patterns.

## 4. Discussion

Many published terrestrial reptile spatial ecology papers reuse the same two methods: Minimum Convex Polygon (MCP), and Kernel Density Estimation (KDE), or variants. Both MCPs and KDEs produced high error rates and failed to properly reflect simulated reptile home ranges. While originally intended for GPS telemetry, we found that dBBMMs perform well across a range of lower fix rates sampling regimes, and for our three archetypical reptile species.

### 4.1. Choice of fix frequency and estimator impacts estimations

The data resampling throughout different tracking regimes led to a 91.7–99.9% data loss from our starting point at 30-minute time steps: removing non-relocations (regime 8) still reduced data points by 70.4–80.0%. Seamen *et al*., (1999) suggested a minimum of 30-50 locations and both regimes 6 (*n* = 52) and 7 (*n* = 12) failed to meet this criteria. A more stringent criteria (Girard *et al.*, 2002) recommending 300 locations also excludes regime 5 (*n* = 104). Based on this fact alone, many reptile studies likely fail to meet KDE requirements.

The use of MCP and KDE *h*_*ref*_ produced large false positive errors, which if carried forward are liable to impact habitat and space-use inferences (Fieberg, 2007; Nilsen *et al.*, 2008). By comparison, both KDE LSCV and dBBMM estimations fared better, although LSCV failed to produce F-measures comparable to dBBMMs under low-resolution tracking regimes. Thus, dBBMMs can improve upon both traditional MCP and fixed KDE methods. As a fix-frequency independent method (Kranstauber *et al.*, 2012), dBBMMs performed most consistently across sampling regimes with the lowest error rates, even in low-resolution datasets. To match dBBMM performance at the sparsest regimes (*n* = 12) KDEs required four times the data. Maximizing performance under low-resolution regimes is critical for VHF studies because the data are time, effort, and cost intensive (Recio *et al.*, 2011).

Furthermore, dBBMMs require no *a priori* knowledge of an animal’s movements (necessary to identify the correct smoothing bandwidth for KDEs), and can be put to use with current telemetry practices or to re-analyse previously collected VHF data. The dBBMM method is easily compatible with low-resolution data from herpetofauna spatial ecology studies still reliant on VHF. As gains from long-term high-resolution tracking methods (GPS) still remain elusive for herpetofauna (Price-Rees, Brown, & Shine, 2013; Wolfe *et al.*, 2018), improving analytic methods represents a cheap, immediate alternative.

At high resolutions the KDE LSCV came closest to performing comparably with dBBMMs despite critical flaws beyond failing the initial point independence assumption. Under higher resolution tracking regimes, the LSCV algorithm fails to converge making the smoothing parameter estimate unusable (supporting findings from Hemson *et al*., 2015). High site fidelity in reptiles leads to unstable KDE LSCV because non-convergence issues are compounded by large numbers of identical locations or very tight clusters (*i.e.* site fidelity). We did not simulate any site fidelity which could inflate LSCV performance. Hemson *et al.*, (2015) suggest ignoring site fidelity in simulation studies leads to inappropriate conclusions advocating KDE LSCV (*e.g.* Worton, 1995; Seaman & Powell, 1996; Seaman *et al.*, 1999). Even with optimal conditions for LSCV, dBBMMs performed similarly or better.

Removing non-relocations (regime 8) improved KDE LSCV while hindering dBBMMs. However, this fix compromises the biological relevance of home range estimates (see De Solla, Bonduriansky, & Brooks, 1999) as the autocorrelated nature of animal movement is inherently biologically relevant (Cushman, Chase & Griffin, 2005). The loss of stationary data points harms inferences drawn upon species that shelter for long periods. Explorations using real GPS data show consistent problems with KDE LSCV omission error, leading to severe undersmoothing, and frequent convergence failures (Hemson *et al*., 2005). Jones, Marron, & Sheather (1996) found that LSCV smoothed utilization distributions had unacceptable variability, that can undermine comparisons between individuals, populations or studies.

Archetypal species movement characteristics influenced our range estimates (MCP, KDE and dBBMM). The active hunter (Species 1), with its sporadic long-distance moves, had lower F-measures and higher error rates than the ambush predators (Species 2 and 3). When comparisons between species are required, researchers should explore how regime and estimation method effect comparisons. Ideally, researchers should be able to access original data from previous studies. We encourage greater use of open data repositories in reptile studies (*e.g.* Movebank). To date, reptile data on Movebank is lacking (11 species, 10 testudines and 1 serpentes). Without readily available data, researchers cannot confidently compare between species.

### 4.2. Caveats

Herpetofauna and VHF tracking studies can be plagued with uncertainty due to inhospitable terrain and associated costs. Failures to detect animals during tracking are inevitable, and we did not assess how the frequency of missed or inconsistent tracks affects each method. Our results indicate that non-symmetrical tracking regimes (*e.g.* tracks performed on Tuesdays and Thursdays) still appear to work well with dBBMMs. Ultimately, accuracy of home range estimation will be dependent on resources, tracking frequency and study duration (Mitchell, White, & Arnold, 2019). All directly impact the viability of answering research questions. A clearly defined research question (Fieberg & Börger, 2012) enables researchers to identify potential trade-offs in context.

While dBBMMs provide a more direct modelling approach for movements –a critical component of assessing habitat use (Van Moorter, Rolandsen, Basille, & Gaillard, 2016)– there is scope for more advanced methods when more is known about a species’ movement patterns. dBBMMs provide an instant option for estimating movement pathways of herpetofauna because they require no *a priori* knowledge. In cases where more data are available, researchers can look at methods that integrate more about the landscape, such as dBBMM with covariates (Kranstauber, 2019), or behaviour (Michelot & Blackwell, 2019). The more advanced methods may require data at higher resolution than is feasibly collectable by VHF.

### 4.3. Recommendations and conclusions

Researchers must consider tracking regime during study design. There are practical considerations of cost, time and ethics, but they must be paired with how the tracking regime will directly impact estimations and, ultimately, the ability to answer research questions. There will always be spatial uncertainty. Tracking regime should minimize spatial uncertainty with reference to the research question and targeted behaviours (Fleming *et al.*, 2014; Schlägel & Lewis, 2016; Bastille-Rousseau *et al.*, 2017). Direct consideration of how biology and movement impact home range will improve inferences drawn from telemetry studies.

The insights into reptile ecology can be invaluable despite data collection costs, and data utility should be maximized. Better home range estimators are an inexpensive way of optimizing returns from tracking data compared to technological advances or increasing field work. Reptile movement is peculiar: we revealed the impact of long-term sheltering (essentially a zero-inflated movement dataset) on home range estimations, which introduced error by under- and over-smoothing with traditional estimators. Inferences based on traditional estimators have likely led to biases in reptile studies. Carrying these biases forward can lead to misallocation of resources.

Our study concurs with previous studies *e.g*. Signer *et al.* (2015) stating problems with both MCP and KDEs. Despite known problems researchers continue to justify use of MCPs and KDEs to maintain comparability with previous studies. We find this deeply flawed in cases where tracking regime or estimator differ which produce dramatically different error rates. However, we also demonstrate the stability of dBBMMs and their suitability for comparisons. The information provided here can help optimise reptile spatial ecology by yielding more accurate and reproducible home range estimations.

## Acknowledgements

We thank Suranaree University of Technology, Institute of Science and Institute of Research and Development for logistic support and facilitating our research. We also thank King Mongkut’s University of Technology Thonburi for support.

