## Supporting Information for "Revisiting reptile home ranges: moving beyond traditional estimators with dynamic Brownian Bridge Movement Models"

### Appendix S1, Supporting Information

Appendix S1 describes the simulation inputs for the hidden Markov models (HMMs). We have defined three archetype species to represent three different groups within reptile movement ecology: **Species 1** corresponds to highly mobile reptiles (active hunters) with long-term shelter sites (*e.g.* monitor lizards, some elapid snakes); **Species 2** represents less mobile reptiles, able to move long distances but with an ambush predator foraging strategy, which still require long-term shelter sites (*e.g.* pythons, crocodiles); finally, **Species 3** represents smaller reptiles species, also ambush predators, with low likelihood of movement and shorter resting periods (*e.g.* viperid snakes, smaller lizards).

All three archetypes have different state-dependent parameters and transition probabilities, but exhibit the same three behaviour states: “sheltering” (state 1), “moving” (state 2), “resting” (state 3). “Sheltering” behaviour corresponds to relatively long periods of immobility, *i.e.* periods of stasis after predation events, ambush or for thermoregulation (King & Duvall 1990, Webb & Shine 1997); “Moving” behaviour corresponds to discrete episodes of movement (species-dependent step lengths) when searching for prey, mates, ambush or shelter sites and retreats (King & Duvall 1990, Madsen & Shine 1993, Secor & Nagy 1994, Webb & Shine 1998) all reflecting, to some degree, habitat selection; “Resting” behaviour occurs only at night and corresponds to short periods of immobility associated with the circadian cycle (but still allowing for some potential shorter movement episodes throughout the night, unlike sheltering). We have created all our archetype species to be diurnal. The emphasis is not on the behaviour itself, but on modelling for home range and general movement patterns. The state-dependent data streams included step length ($l_t$) and turning angle ($\theta_t$), which were generated from Gamma) and von Mises (α, κ) distributions, respectively. The state-dependent data streams included step length ($l_t$) and turning angle ($\theta_t$), which were generated from Gamma and von Mises distributions, respectively:

$$l_t \mid S_t = s \sim gamma(\mu_s, \sigma_s), \quad \mu > 0,\ \sigma > 0 \quad (1)$$
$$\theta_t \mid S_t = s \sim von\,Mises(\alpha_s, \kappa_s), \ \alpha \,\epsilon\, (\pi, -\pi), \quad \kappa > 0 \quad (2)$$

We have modelled step length for state 2, “Moving” ($\mu_t^l$), as functions of the habitat value of each cell ($h_t$). This allows the simulated animal to showcase a pseudo-preference behavior, moving faster through bad habitats (*i.e.* lower $h_t$) and slower through better habitats (*i.e.* higher $h_t$):

$$\mu_t^l = exp(\beta_0^l + \beta_1^l \times h_t) \quad (3)$$

The von Mises distribution has its mean centred around zero ($\alpha \sim 0$), so there is persistence in the movement direction.

The transitional probability matrices, $\Gamma^{(t)}$, for all species follow the same format where each element $\gamma_{i,j}$ is the probability of switching from state *i* at time *t* to state *j* at time *t*+1 for *i*,*j* ∈ {1=“sheltering”, 2=“moving”, 3=“resting”}:

$$\Gamma^{(t)} = \left(\gamma_{i,j}^{(t)}\right) = \begin{pmatrix} \gamma_{1,2} & \cdots & \gamma_{3,j} \\ \vdots & \ddots & \vdots \\ \gamma_{1,2} & \cdots & \gamma_{3,j} \end{pmatrix} \tag{4}$$

Individuals cannot alternate between states 1 (“sheltering”) and 3 (“resting”) as we believe it is not biologically reasonable to go into sheltering or resting without first going through state 2 (“moving”), *i.e.* without first exhibiting some foraging behaviour. This is reflected by a large negative number (-100), following recommendations from McClintock & Michelot (2018). We also modelled the transition probabilities to vary through time, with the transition of 1→2 and 2→1 being modelled as a *cosinor* function varying monthly (with slight changes by species). The transition of 2→3 and 3→2 we modelled as a *cosinor* function every 24 hours (daily; equal in all three species). This allows us to simulate patterns of both short-term resting (circadian rhythm; $\gamma_{1,2}$) and variable-length sheltering ($\gamma_{2,3}$). For the latter, $\tau_i$ is species-dependent to reflect different sheltering patterns:

$$\gamma_{1,2}^{(t)} = \gamma_{2,1}^{(t)} = [\ cos(2\pi \times t/\tau_i),\ sin(2\pi \times t/\tau_i)\ ],\ \ t \in (0,1,\ldots,23), \tau_i = 24 \tag{5}$$
$$\gamma_{2,3}^{(t)} = \gamma_{3,2}^{(t)} = [\ cos(2\pi \times t/\tau_i),\ sin(2\pi \times t/\tau_i)\ ],\ \ t \in (0,1,\ldots,365) \tag{6}$$

The low number of reptile studies using GPS technology, and inconsistent data accessibility, limited our ability produce literature-derived *a priori* estimations of step length and turning angles. Therefore, we relied on the results of exploratory data analyses performed on simulations for each of our archetype species. By testing multiple parameters, we were able to identify those parameters that produced datasets approximating reality: Reality as inferred by the summary statistics provided in reptile telemetry studies.

### Species 1

#### “*Active hunter, long-distance moves, long-term sheltering*”

Highly mobile reptile species are usually active hunters (*i.e.* forage by actively searching for prey), as they show higher values of standard metabolic rates than ambush predators (Stuginski *et al.*, 2018). Foraging activity can vary widely among active hunting snake species, depending on the kind of prey hunted (Stuginski *et al.*, 2018), we chose to reflect a specialization on larger prey as it may be favoured over small prey items (*e.g.* Purwandana *et al.*, 2016; Wallace & Leslie, 2008). As such, this will lead to a longer use of sheltering sites (for thermoregulation and digestion of prey items).

| Data stream | State 1 - Sheltering | State 2 - Moving | State 3 - Resting |
|---|---|---|---|
| Step length | $s_1 \sim gamma$ (0.01, 0.01) | $s_2 \sim gamma$ (20, 10) | $s_3 \sim gamma$ (0.2, 0.2) |
| Turning angle | $s_1 \sim von\ Mises$ (0, 0.01) | $s_2 \sim von\ Mises$ (0, 0.01) | $s_3 \sim von\ Mises$ (0, 0.01) |

The transition probability matrix for Species 1 is the following:

$$\Gamma = \begin{pmatrix} -5 & -100 & -10 & 0 & -100 & 0 \\ 6 & 0 & -4 & 0 & 0 & 0 \\ -6 & 0 & 5 & 0 & 0 & 0 \\ 0 & 0 & 0 & 4 & 0 & -4 \\ 0 & 0 & 0 & 2 & 0 & -2 \end{pmatrix}$$

We set the transition probabilities between states 1→2 and 2→1 as a *cosinor* function fluctuating every 21 days ($\tau_i$), simulating a pattern of long-term sheltering (~2-3 weeks).

### Species 2

#### "*Ambush predator, long-distance moves, long-term sheltering*"

Sedentary, "sit-and-wait" ambush predators include many snake species, such as vipers, pythons, boas, as well as some colubrids and elapids (Shine, 1980; Stuginski *et al.*, 2018). Snakes with an ambush foraging strategy consume a wide range of meal sizes (Glaudas *et al.*, 2019): We chose to model Species 2 after a specialization on larger prey. The main component of an ambush predator is spending long periods in the same location (Reinert *et al.*, 1984; Webb & Shine, 1997). "Sit-and-wait" predators also usually feed less frequently than active hunters (Huey & Pianka, 1981).

| Data stream | Sheltering | Moving | Resting |
|---|---|---|---|
| Step length | $s_1 \sim gamma$ (0.01, 0.01) | $s_2 \sim gamma$ (15, 5) | $s_3 \sim gamma$ (0.2, 0.2) |
| Turning angle | $s_1 \sim von\ Mises$ (0, 0.01) | $s_2 \sim von\ Mises$ (0, 0.01) | $s_3 \sim von\ Mises$ (0, 0.01) |

The transition probability matrix for Species 2 is the following:

$$\Gamma = \begin{pmatrix} -13 & -100 & -11 & 0 & -100 & 0 \\ 5.5 & 0 & -4 & 0 & 0 & 0 \\ -5.5 & 0 & 5 & 0 & 0 & 0 \\ 0 & 0 & 0 & 4 & 0 & -4 \\ 0 & 0 & 0 & 2 & 0 & -2 \end{pmatrix}$$

We set the transition probabilities between states 1→2 and 2→1 as a *cosinor* function fluctuating every 21 days ($\tau_i$), simulating a pattern of long-term sheltering (~2-3 weeks).

### Species 3

#### "*Ambush predator, short-distance moves, short-term sheltering*"

In contrast to Species 2, we chose to model Species 3 with a smaller body mass, reflected by short-distance moves and specialization on smaller prey (resulting in shorter sheltering events, as digestion is faster with smaller prey items). In particular, species 3 behaves similarly to pit vipers (family Viperidae): ambush predators, low energetic requirements, infrequent feeding events, as well as reduced movement rate and home range size in comparison to other larger snake species (Mushinsky, 1987; Macartney *et al.*, 1988).

| Data stream | Sheltering | Moving | Resting |
|---|---|---|---|

| | | | |
|---|---|---|---|
| Step length | $s_1$ ~ Gamma(0.01, 0.01) | $s_2$ ~ Gamma(2, 5) | $s_3$ ~ Gamma(0.5, 0.5) |
| Turning angle | $s_1$ ~ von Mises(0, 0.01) | $s_2$ ~ von Mises(0, 0.01) | $s_3$ ~ von Mises(0, 0.01) |

The transition probability matrix for Species 3 is the following:

$$\Gamma = \begin{pmatrix} -5 & -100 & -5 & 0 & -100 & 0 \\ 2 & 0 & -2 & 0 & 0 & 0 \\ -2 & 0 & 2 & 0 & 0 & 0 \\ 0 & 0 & 0 & 4 & 0 & -4 \\ 0 & 0 & 0 & 2 & 0 & -2 \end{pmatrix}$$

We set the transition probabilities between states 1→2 and 2→1 as a *cosinor* function fluctuating every 7 days ($\tau_i$), simulating a pattern of short-term sheltering (~1 week), in opposition of the previous two species (Siers *et al.*, 2018).

### Appendix S2, Supporting Information

In this paper, we were interested in evaluating how regime and method affected the F-measures of the three archetype reptile species. We ran Bayesian mixed effects models using the *brms* R package (Bürkner, 2017), as to estimate the expected values and the uncertainty associated with each parameter. We built three model sets (one for each species), and each model was run with six independent Markov chains, with 6000 iterations each, discarding the first 1000 iterations per chain as burn-in.

In order to account for the individual variation, we included individual (IND) as a random effect. The variation of F-measures between different methods may not identical for each individual; to address this issue, we added a varying slope for the effect of method (1 + METHOD | IND). To eliminate the number of divergent transitions that could cause bias in the posterior samples, we increased the adapt delta from 0.8 (default) to 0.99.

We tested three different models for each species: 1) Model 1, *null model*, with intercept only; 2) Model 2, with method and regime as group effects; and 3) Model 3, with method and regime as group effects and an added interaction effect.

```
brm1 <- brm(fmeasure ~ 1 + (1 + METHOD|IND),
        family = "beta", control = list(adapt_delta = .99),
             chains = 6, iter = 6000, warmup = 1000)

brm2 <- brm(fmeasure ~ METHOD + REG + (1 + METHOD|IND),
        family = "beta", control = list(adapt_delta = .99),
             chains = 6, iter = 6000, warmup = 1000)

brm3 <- brm(fmeasure ~ METHOD * REG + (1 + METHOD|IND),
        family = "beta", control = list(adapt_delta = .99),
             chains = 6, iter = 6000, warmup = 1000)
```

For model selection, we used the leave-one-outcross-validation with Pareto smoothed importance sampling (PSIS-LOO CV; Vehtari *et al.*, 2016) This method is implemented with the function `loo()`, calculating the expected log pointwise predictive density (ELPD), providing an estimate of a model's predictive accuracy. It is a better evaluation method than simpler estimates like AIC (Akaike information criterion) and DIC (deviance information criterion). The diagnostic used to determine whether a point is influential is the pareto-k ($\hat{k}$) parameter, with any $\hat{k}$ value above 0.7 considered problematic. To address this issue, we refitted all models by retaining each influential observation at a time (*i.e.* `loo()` function with `reloo = TRUE`).

We then conducted model validation by posterior predictive check using function `pp_check()`, to show how close our simulated samples were to the observed data. Finally, we calculated the effect size (Bayesian $R^2$; Gelman *et al.*, 2018).

#### Species 1

##### “*Active hunter, long-distance moves, long-term sheltering*”

An initial run for species 1 revealed model 2 with one observation with a $\hat{k} > 0.7$, and model 3 with two such observations. After re-running the leave-one-outcross-validation, all $\hat{k}$ estimates are $< 0.7$; these are the final model comparisons:

| Model | Formula | LOOIC | SE | ΔELPD | ΔSE |
|---|---|---|---|---|---|
| `brm3` | `METHOD * REG` | -3262.4 | 104.0 | 0 | 0 |
| `brm2` | `METHOD + REG` | -2024.3 | 67.9 | -619.1 | 37.6 |
| `brm1` | *Null model* | -1643.9 | 71.0 | -809.3 | 36.9 |

Model 3 (METHOD * REG) had the lowest LOOIC values, indicating the highest predictive accuracy. In addition, all parameters have $\hat{R} = 1.00$ with reasonably large effective sample sizes, strongly suggesting that the sampling algorithm has converged to the posterior distribution. The distribution of the observed values versus simulated values also suggest that the model fitted is doing reasonably well (Figure S1). The visual inspection of the traceplots also confirmed chain convergence. The effect size of model 3 (Bayesian $R^2$) was 0.960 ± 0.001 (CrI: 0.958–0.962).

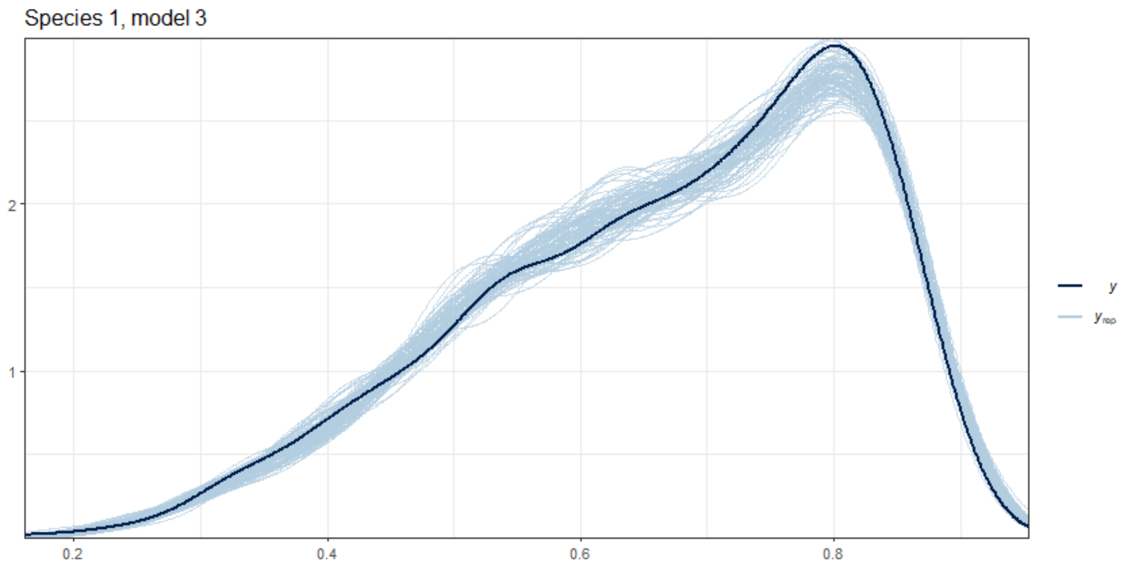


Figure S1. Distribution of observed values, compared with distributions of simulated values drawn from the posterior predictive distribution, using Species 1 Model 3, and only showing the first 100 samples for ease of visualization.

### Species 2

*"Ambush predator, long-distance moves, long-term sheltering"*

An initial run for species 2 revealed model 3 with five observations with a $\hat{k} > 0.7$. After re-running the leave-one-outcross-validation, all $\hat{k}$ estimates are $< 0.7$; these are the final model comparisons:

| Model | Formula | LOOIC | SE | ΔELPD | ΔSE |
|---|---|---|---|---|---|
| `brm3` | `METHOD * REG` | -3064.7 | 117.4 | 0 | 0 |
| `brm2` | `METHOD + REG` | -2047.0 | 73.8 | -510.8 | 48.3 |
| `brm1` | *Null model* | -1765.3 | 80.9 | -649.7 | 47.9 |

Model 3 (METHOD * REG) had the lowest LOOIC values, indicating the highest predictive accuracy. In addition, all parameters have $\hat{R} = 1.00$ with reasonably large effective sample sizes, strongly suggesting that the sampling algorithm has converged to the posterior distribution. The distribution of the observed values versus simulated values also suggest that the model fitted is doing reasonably well (Figure S2). The visual inspection of the traceplots also confirmed chain convergence. The effect size of model 3 (Bayesian $R^2$) was 0.946 ± 0.002 (CrI: 0.942–0.949).

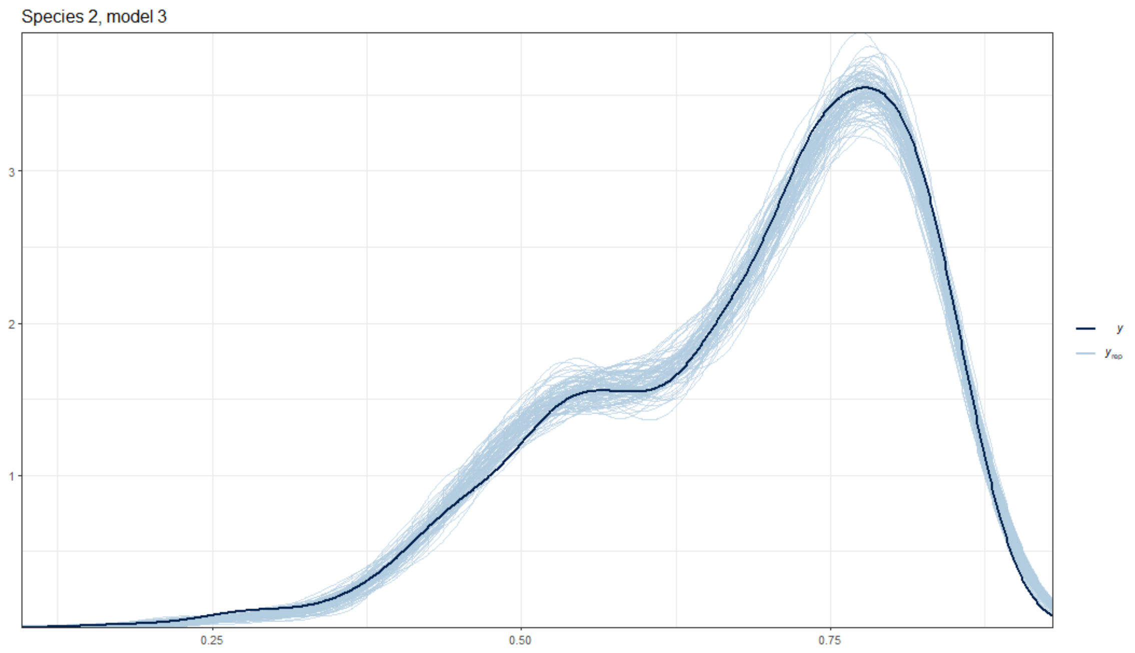


Figure S2. Distribution of observed values, compared with distributions of simulated values drawn from the posterior predictive distribution, using Species 2 Model 3, and only showing the first 100 samples for ease of visualization.

### Species 3

#### "*Ambush predator, short-distance moves, short-term sheltering*"

An initial run for species 3 revealed model 1 with one observation with a $\hat{k} > 0.7$, and model 3 with 9 such observations. After re-running the leave-one-outcross-validation, all $\hat{k}$ estimates are $< 0.7$; these are the final model comparisons:

| Model | Formula | LOOIC | SE | ΔELPD | ΔSE |
|---|---|---|---|---|---|
| `brm3` | `METHOD * REG` | -2956.0 | 81.1 | 0.0 | 0.0 |
| `brm2` | `METHOD + REG` | -2274.5 | 57.3 | -340.8 | 36.3 |
| `brm1` | *Null model* | -1967.6 | 66.5 | -494.1 | 40.5 |

Model 3 (METHOD * REG) had the lowest LOOIC values, indicating the highest predictive accuracy. In addition, all parameters have $\hat{R} = 1.00$ with reasonably large effective sample sizes, strongly suggesting that the sampling algorithm has converged to the posterior distribution. The distribution of the observed values versus simulated values also suggest that the model fitted is doing reasonably well (Figure S2). The visual inspection of the traceplots also confirmed chain convergence. The effect size of model 3 (Bayesian $R^2$) was 0.905 ± 0.004 (CrI: 0.897–0.911).

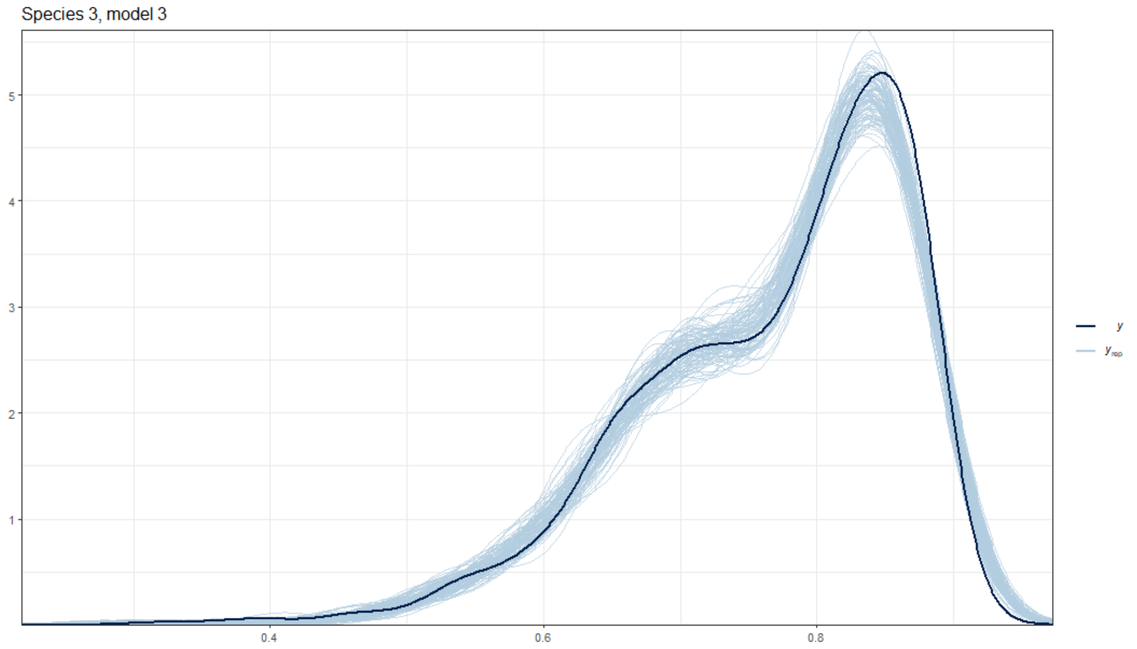


Figure S3. Distribution of observed values, compared with distributions of simulated values drawn from the posterior predictive distribution, using Species 3 Model 3, and only showing the first 100 samples for ease of visualization.
